# CryoFAST: A Fab-Aided Cryo-EM Workflow for High-Resolution Structure Determination of Sub-50-kDa Protein Targets

**DOI:** 10.64898/2026.10.07.757472

**Authors:** Xiaorun Li, Sean Deng, Weihua Li, Yue Qian, Yawen Li, Juan Du, Le Li, Dongming Qian

## Abstract

Single-particle cryo-electron microscopy (cryo-EM) plays an increasingly important role in structure-based drug discovery (SBDD) by revealing target protein-ligand interactions at the molecular level, but its application has largely focused on large- and medium-sized protein complexes. Many therapeutically important targets are smaller than 50 kDa and remain highly challenging for cryo-EM analysis because of low signal-to-noise ratios and difficulties in particle alignment. Here, we describe cryoFAST, an integrated workflow that couples ligand-compatible Fab discovery and structure-guided computational maturation with biochemical triage, fiducial-complex assembly, and single-particle cryo-EM. Through phage-display screening and/or AI-guided design, we developed high-affinity Fab binders of approximately 50 kDa that stabilize the target without disrupting ligand-target interactions and then assembled Fab-target complexes for cryo-EM analysis. Using the therapeutically relevant extracellular domain (∼20 kDa) of interleukin-4 receptor α (IL-4Rα) as a proof of concept, we determined its cryo-EM structure at 2.74 Å resolution and established a framework for elucidating ligand-binding modes. This workflow provides a practical, efficient, and potentially broadly applicable route to the structural characterization of small protein targets that are otherwise challenging to study. By extending the applicability of cryo-EM to high-value but technically challenging small targets, cryoFAST may open new opportunities for structure-based drug discovery.

## Introduction

Single-particle cryo-EM can now provide near-atomic structures for a broad range of macromolecular assemblies and has become increasingly integrated into structure-based drug discovery (SBDD) [1–5]. Unlike crystallography, cryo-EM does not require formation of an ordered crystal lattice and can therefore be used to analyze conformationally heterogeneous complexes and ligand-bound states under frozen-hydrated conditions [6,7]. These features have made cryo-EM particularly effective for large protein complexes and membrane proteins and increasingly useful for defining target-ligand and target-antibody interactions relevant to drug discovery [4,5,8,9].

The performance of single-particle analysis nevertheless remains strongly dependent on particle size, rigidity, and the presence of recognizable features [10]. At a fixed electron dose, the signal contributed by a particle decreases with molecular mass, whereas shot noise and the background from vitreous ice remain substantial. This compromises particle detection and angular assignment and can amplify the effects of conformational flexibility. Proteins below 50 kDa are consequently underrepresented among high-resolution cryo-EM structures, even though many therapeutically relevant enzymes, extracellular domains, cytokines, and protein-protein interaction modules fall within this size range [11–13].

Several strategies have been developed to overcome this size barrier. These include fusion to larger domains, attachment to symmetric scaffolds, and binding of Fabs, nanobodies, or engineered binder-scaffold modules [11–19]. A fiducial can increase particle mass, enlarge the maximum particle dimension, provide asymmetric features for alignment, and sometimes stabilize a flexible target. However, existing approaches often require genetic modification of the target, optimization of fusion geometry, or access to a pre-existing binder. For SBDD applications, the fiducial must satisfy an additional constraint: it should not occlude the ligand-binding site or distort the target conformation intended for analysis. In practice, no single convenient and robust approach is yet widely used for cryo-EM-enabled SBDD of small therapeutic targets, which therefore remain challenging for the field.

We reasoned that small-target cryo-EM would benefit from an end-to-end workflow in which binder discovery, epitope selection, affinity and stability optimization, ligand-compatibility testing, and cryo-EM complex assembly are treated as connected design steps rather than independent activities. We therefore developed cryoFAST, a **cryo**-EM-enabled **F**ab-**A**ided **S**tructure determination workflow for challenging sub-50 kD **T**herapeutic protein targets, centered on the following practical criteria: (i) recognition of the unmodified target; (ii) an epitope separated from the intended ligand-binding region; (iii) high occupancy and biochemical homogeneity; and (iv) a sufficiently rigid target-Fab geometry for particle alignment. The workflow uses phage display [20] to obtain ligand-compatible target-binding antibody fragments, applies AI-enabled *in silico* sequence design to improve binding affinity, and advances biochemically stable complexes to cryo-EM. The intended output is not simply a larger particle, but a structurally interpretable assembly in which the pharmacological binding site remains readily accessible.

As a proof of concept, we applied cryoFAST to the human IL-4Rα extracellular domain (ECD). IL-4Rα is a central component of type I and type II IL-4/IL-13 receptor complexes and is a clinically validated extracellular target [21,22]. Although antibody therapeutics targeting this pathway are clinically established and multiple structures of IL-4Rα-containing complexes are available [21,23,24], structural information for soluble interleukin-receptor ECDs in complex with small-molecule or peptide ligands remains comparatively limited. Their modest molecular size, glycosylation, and conformational flexibility create practical challenges for both crystallography and cryo-EM, making IL-4Rα a useful test of whether a ligand-compatible Fab workflow can generate a high-resolution cryo-EM target structure. Here, we describe the complete workflow, present a 2.74-Å IL-4Rα ECD fiducial-complex reconstruction, and define both the practical advantages and current limitations of the approach.

## Results

### Workflow of cryoFAST

The primary objective of cryoFAST is to convert a small target into a larger, more readily alignable particle while preserving the native structural state and ligand-binding surface of the target. The workflow comprises three experimental phases (Fig. 1). In Step 1, target-specific binders are generated through phage display and/or AI-guided design, with explicit selection for epitopes compatible with the intended ligand-binding site. In Step 2, candidate Fabs are evaluated for target binding, ligand compatibility, and formation of stable and homogeneous complexes. In Step 3, the selected Fab-target complex is assembled and subjected to cryo-EM grid optimization, data collection, image processing, and structural validation.

**Figure 1.**
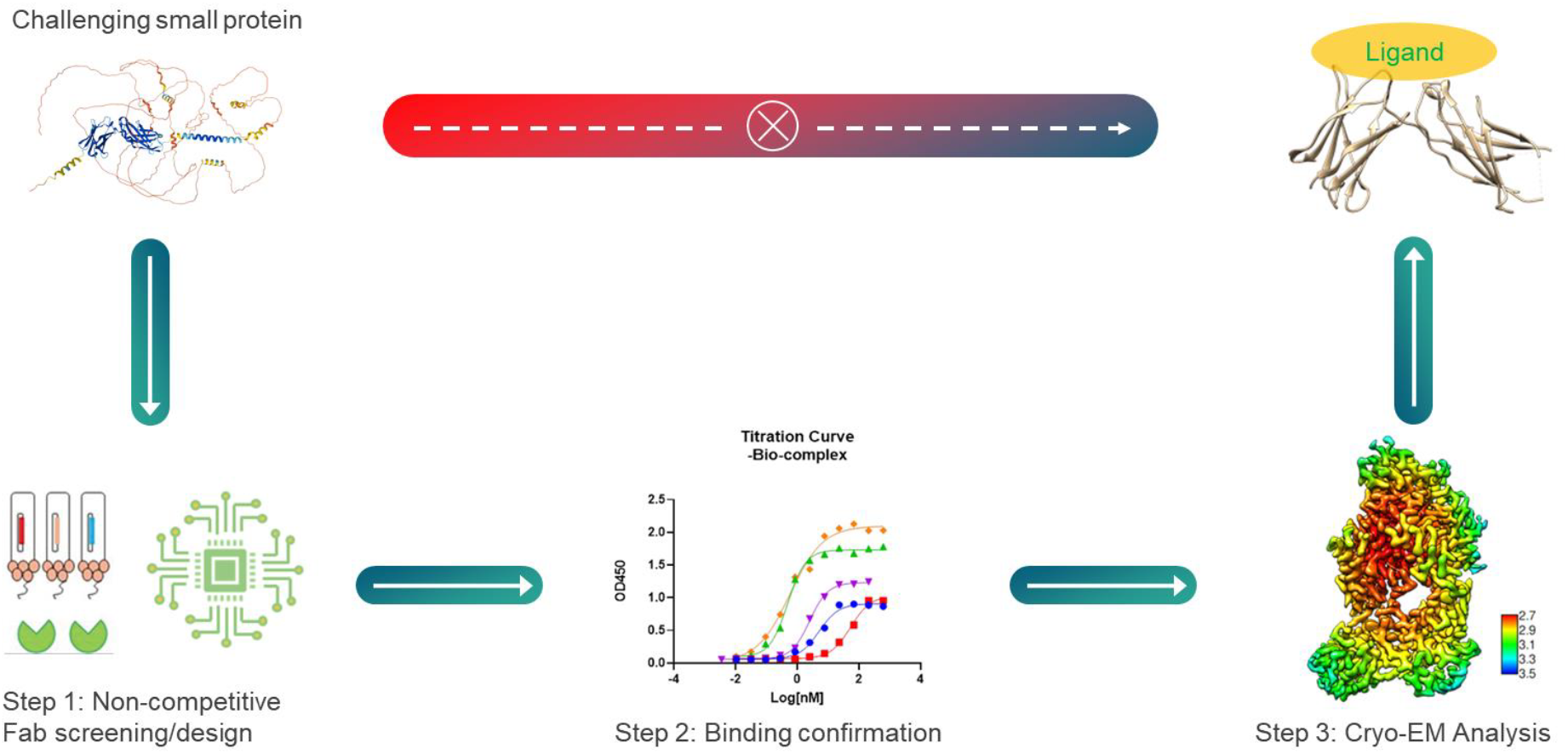
Overview of the cryoFAST workflow. **Step 1:** Non-competitive Fabs that preserve ligand binding are discovered through phage display screening and/or AI-driven design. **Step 2:** The optimal Fab is selected by biophysical/biochemical triage. **Step 3:** The Fab-target complex is assembled for cryo-EM structure determination, enabling elucidation of target-ligand interactions at atomic resolution.

**Figure 2.**
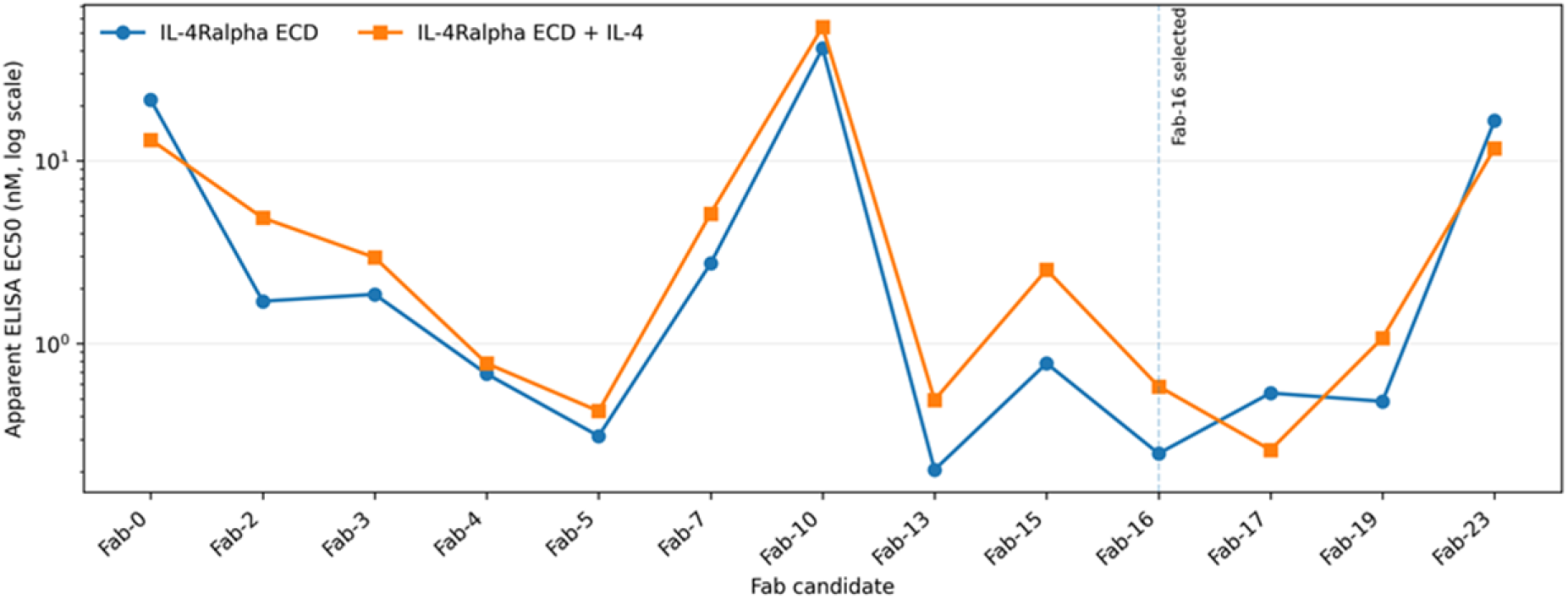
Apparent binding potency of Fab candidates. Apparent EC50 values derived from concentration-response ELISA are shown for binding to the IL-4Rα ECD and a preassembled IL-4Rα-IL-4 complex.

The workflow is modular at the binder-discovery stage but uses common decision gates before cryo-EM. A candidate advances only when it shows high target occupancy, retains binding to the ligand-loaded target, forms a discrete complex by size-exclusion chromatography (SEC), and produces well-resolved two-dimensional (2D) classes with multiple views. These gates are intended to prevent a common failure mode in fiducial-assisted cryo-EM: increasing nominal molecular mass without creating a rigid and homogeneous imaging particle. Below, we use IL-4Rα as an example to illustrate the workflow.

### Phage-display screening identifies an IL-4Rα-binding starting clone

Human IL-4Rα ECD was used as the antigen for selection from VIVA Biotech’s in-house human single-chain variable fragment (scFv) phage library. The antigen was immobilized at 5 µg ml^−1^, and two rounds of panning were performed with triethylamine elution and TG1 amplification. Individual clones were screened by phage ELISA against IL-4Rα and a preassembled IL-4Rα-IL-4 complex. Binding to IL-4 alone was included as a specificity control. Clones that recognized both receptor preparations but not IL-4 were prioritized, thereby enriching for epitopes that remained accessible after orthosteric cytokine binding.

A starting clone, designated scFv-IL-4R-initial, was selected for sequence analysis and conversion into Fab format. This clone provided a target-binding scaffold but showed only modest apparent binding potency in the subsequent concentration-response ELISA, motivating optimization guided by AI-based structure prediction.

### Structure-prediction-guided optimization generates additional Fab candidates

A structural model of the starting Fab bound to IL-4Rα was generated using AlphaFold2-Multimer or an equivalent multimeric structure-prediction workflow [25,26]. The model placed the Fab on the D2 region of IL-4Rα, away from the expected IL-4-binding surface. A 100-ns molecular-dynamics simulation was then used to rank interfacial contacts, including hydrogen bonds, salt bridges, and hydrophobic interactions. Interface-focused saturation mutagenesis was performed *in silico*, with emphasis on VH CDR3 and VL CDR3 positions. Candidate substitutions were ranked using an in-house scoring matrix and manual structural review, yielding 23 designed Fab variants.

### Triage of candidate Fabs with strong target binding and retained ligand compatibility

The designed variants and the starting Fab were expressed and screened against both IL-4Rα and IL-4Rα-IL-4. Thirteen variants were advanced to concentration-response analysis. The ELISA curves indicate substantial shifts in apparent EC50 relative to the starting Fab (Fab-0), while several optimized variants retained binding to the cytokine-bound receptor (Supplementary Fig. 1 and Supplementary Table 1). Fab-16 showed apparent EC50 values of approximately 0.25 nM for IL-4Rα and 0.58 nM for IL-4Rα-IL-4, compared with 21.5 nM and 13.0 nM, respectively, for the starting Fab. Because these values were derived from ELISA concentration-response curves, they are reported here as apparent EC50 values rather than equilibrium dissociation constants.

Fab-16 was selected for structural studies based on its combined binding profile, ligand compatibility, biochemical behavior, and initial cryo-EM screening results. The final selection triage criteria should incorporate expression yield, SEC behavior, and kinetic affinity if these measurements are available.

### Cryo-EM analysis yields a 2.74-Å reconstruction of the IL-4Rα fiducial complex

The nominal mass of the assembly is expected to be substantially larger than that of IL-4Rα alone, while the elongated Fab and the additional anti Fab nanobody provides recognizable shape features for image alignment. Importantly, the use of a non-blocking Fab allows a single fiducial assembly to be prepared before addition of an orthosteric ligand, creating a potential route to repeated ligand-soaking or co-incubation experiments.

Cryo-EM grids were prepared and then screened using a Glacios microscope. Data were collected on a Titan Krios G4 equipped with a Falcon 4i direct electron detector, leading to the final reconstruction at an overall resolution of 2.74 Å according to the gold-standard FSC = 0.143 criterion (Fig. 3A, B). The local-resolution map (Fig. 3B) spans approximately 2.7–3.5 Å, with sufficient density for side-chain interpretation across the IL-4Rα-Fab interface.

**Figure 3.**
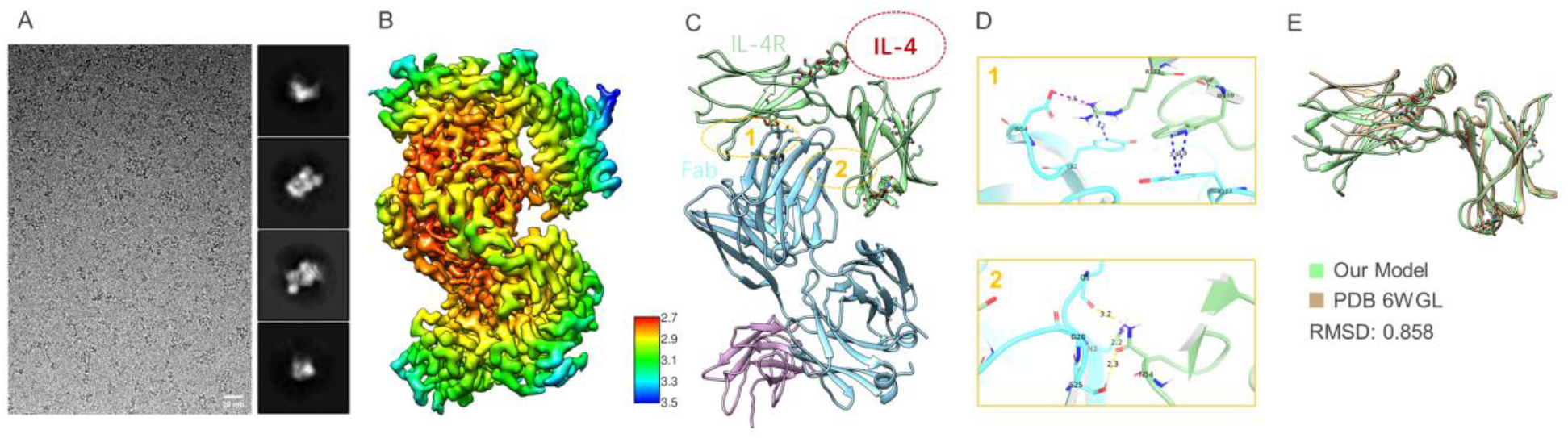
Proof-of-concept validation of cryoFAST using the IL-4Rα ECD (∼20 kDa). **A**, Representative cryo-EM micrograph and 2D class averages of the IL-4Rα ECD/Fab-16/anti-Fab nanobody complex. **B**, 3D reconstruction of the IL-4Rα ECD/Fab-16/anti-Fab nanobody complex, color-coded according to local resolution. **C**, Color-coded ribbon diagram of the IL-4Rα ECD/Fab-16/anti-Fab nanobody complex atomic model. The orthosteric IL-4-binding site is indicated by a red circle. **D**, Enlarged views of the two IL-4Rα/Fab-16 interfaces, with key interactions highlighted. **E**, Structural superposition of the IL-4Rα ECD/Fab-16 model with the previously reported IL-4Rα structure (PDB 6WGL). The close alignment between the two models, with a backbone RMSD of 0.858 Å, indicates that Fab binding does not substantially alter the IL-4Rα conformation and is therefore unlikely to interfere with orthosteric ligand binding.

### The Fab recognizes an epitope distal from the IL-4-binding site and does not perturb the receptor backbone

An atomic model was built into the 2.74-Å map using an IL-4Rα crystal structure as the initial receptor template [23]. The final model contains the IL-4Rα ECD, Fab-16, and the anti-Fab nanobody (Fig. 3C). The Fab contacts IL-4Rα through two principal interfaces, including hydrogen bonding between IL-4Rα N54 and Fab heavy-chain residues N3 and S25, a π-π interaction between IL-4Rα W139 and Fab heavy-chain Y107, and combined cation-π and salt-bridge contacts between IL-4Rα R173 and Fab heavy-chain Y52 and D54, respectively (Fig. 3D).

The Fab epitope is spatially separated from the orthosteric IL-4-binding region. Consistent with this geometry, Fab-16 retained ELISA binding to the preassembled IL-4Rα-IL-4 complex. Superposition of the IL-4Rα chain with PDB 6WGL [23] yielded a backbone RMSD of 0.858 Å, supporting close global agreement between the receptor conformations (Fig. 3E). This comparison indicates that Fab binding does not substantially alter the overall IL-4Rα fold, although direct ligand-bound biophysical/structural data will be required to demonstrate preservation of specific small-molecule or peptide binding modes.

### Generality beyond IL-4Rα

We have applied a similar workflow to multiple additional small therapeutic targets and obtained high-resolution cryo-EM structures that supported early drug-discovery programs. Because these projects are subject to confidentiality and intellectual-property restrictions, further target-specific details cannot be disclosed here. Accordingly, the IL-4Rα study remains the only fully disclosed example in this manuscript.

## Discussion

This study establishes a proof-of-concept workflow for extending single-particle cryo-EM to a small extracellular receptor domain. The central feature of cryoFAST is not simply the addition of molecular mass. Instead, the workflow links binder discovery, epitope selection, AI-guided optimization, ligand-compatibility screening, and cryo-EM validation. In the IL-4Rα example, this strategy converted an approximately 20-kDa target into a > 70-kDa assembly and enabled an atomic resolution cryo-EM reconstruction.

Fab-assisted cryo-EM has a long history, and several modular scaffolds have been developed to improve imaging of small targets [11,15,18]. In SBDD, the principal objective is often to define target-ligand interactions rather than the apo structure alone. cryoFAST complements these approaches by placing particular emphasis on ligand compatibility. It avoids genetic modification of the target and prioritizes selection of a target-specific epitope that remains accessible in the ligand-bound state. This approach may be especially useful for soluble extracellular domains that are difficult to crystallize reproducibly or need to be analyzed with multiple ligand chemotypes.

Recent applications of the workflow have included both commercially available antibodies and in-house-developed antibodies that were screened and selected for ligand compatibility, leading to successful structure determination of multiple small protein targets. These datasets further informed the development and refinement of the workflow, although they are not presented here as formal cross-target validation because of confidentiality restrictions. In addition, with recent breakthroughs and rapid advances in de novo antibody design [27–29], we anticipate that the design of antibodies against defined antigens with selective epitope targeting will become increasingly accessible. This should further expand the applicability of cryoFAST by enabling more efficient generation of ligand-compatible fiducial binders tailored to individual targets.

We acknowledge several limitations of the current cryoFAST workflow. First, the method is demonstrated in detail for a single target, and its broader generality remains to be established using disclosed cross-target datasets. Based on our experience and discussions with other researchers, developing suitable Fabs can be challenging for certain target classes, such as small membrane proteins. Second, the current approach may be less effective for very small targets (<10 kDa), even in the presence of a Fab, and successful implementation may require substantial optimization and empirical screening. Third, ligand series that bind to distinct sites may require Fabs recognizing different epitopes. Fourth, continued advances in cryo-EM hardware and image-processing methods, including laser phase plates, may ultimately enable direct high-resolution analysis of progressively smaller targets and reduce the need for fiducial markers [30]. Despite these limitations, cryoFAST provides a practical development framework and a clear experimental roadmap for extending cryo-EM analysis to challenging small protein targets.

## Methods

### IL-4Rα ECD construct, expression, and purification

VIVA Biotech’s in-house, off-the-shelf IL-4Rα ECD and IL-4 proteins were used throughout this study. In brief, an IL-4Rα(26–232)-Avi construct was generated and expressed in a mammalian expression system. The culture supernatant was purified by nickel-affinity chromatography followed by size-exclusion chromatography (SEC). Final protein quality was assessed by SDS-PAGE and western blotting.

### Phage-display selection

IL-4Rα ECD was diluted in phosphate-buffered saline (PBS) to 5 µg ml^−1^ in a total volume of 0.5 ml and immobilized in an immunotube overnight at 4 °C. The tube was washed three times with 5 ml PBS and blocked for 1 h at room temperature with 5 ml 3% (w/v) non-fat milk in PBS. A 100-µl aliquot of a human scFv phage library was mixed with 400 µl blocking solution for 1 h at room temperature and then incubated in the antigen-coated tube for 1 h.

The tube was washed five times with PBS containing 0.1% Tween-20 (PBST) and five times with PBS. Bound phage were eluted with 0.5 ml freshly prepared 0.1 M triethylamine for 15 min and neutralized with 250 µl 1 M Tris-HCl, pH 7.4. Eluted phages were amplified in exponentially growing TG1 cells and rescued with helper phage. Two rounds of panning were performed. Individual colonies were picked into 96-well plates, amplified, and screened by phage ELISA.

### Phage ELISA and clone selection

High-binding, flat-bottom 96-well ELISA plates were coated overnight at 4 °C with IL-4Rα, IL-4Rα-IL-4 complex, or IL-4 at 2 µg ml^−1^ in PBS (100 µl per well). Plates were washed three times with PBS and blocked with 200 µl per well of 3% non-fat milk in PBS for 1 h at room temperature. Phage-containing supernatants were diluted 1:3 in blocking buffer, incubated for 1 h, and detected with M13-HRP (Sino Biological, catalogue no. 11973-MM05T-H, 0.3 µg ml^−1^). Plates were washed three times with PBST (0.1% Tween in PBS) and then three times with PBS, using 300 µl per well. After washing, TMB substrate was added for 15 min, the reaction was stopped with 2 M sulfuric acid, and absorbance was measured at 450 nm. Positive clones were sequenced, and the leading clone was converted to Fab format.

### Structural modeling and computational Fab optimization

The starting Fab and IL-4Rα sequences were submitted to AlphaFold2-Multimer or an equivalent multimeric structure-prediction workflow to generate a model of the complex. The predicted interface was evaluated by molecular dynamics using AMBER for 100 ns. The AMBER ff14SB force field and TIP3P water model were used. The system was solvated in a cubic box with a minimum solute-boundary distance of 10 Å. Following energy minimization, the protein system was heated to 300 K over 5,000 steps and maintained at 300 K for an additional 5,000 steps. The system was then equilibrated for 400 ps. Hydrogen bonds, salt bridges, hydrophobic contacts, and residue-wise interaction contributions were analyzed across the trajectory. Selected CDR positions, particularly in VH CDR3 and VL CDR3, were subjected to in silico saturation mutagenesis. Variants were ranked by calculated interface score, predicted complementarity, formation of new polar interactions, and manual assessment of developability. Twenty-three designs were selected for experimental testing.

### Fab expression and purification

The 23 designed variants and the starting sequence were cloned into Fab expression vectors and transiently expressed in Expi293F cells using the ExpiFectamine 293 system. Total plasmid DNA was used at 1.0 µg per milliliter of transfection volume, and cultures were harvested 5 d after transfection. Culture supernatants were clarified by centrifugation at 2,000 rpm for 5 min followed by 10,000 rpm for 20 min. The supplied patent describes capture on MabSelectSure resin equilibrated in PBS, elution with 50 mM acetate at pH 3.0, immediate neutralization with 1 M Tris at pH 8.0, and buffer exchange into PBS.

### Concentration-response ELISA

Plates were coated with IL-4Rα or IL-4Rα-IL-4 complex at 2 µg ml−1 in PBS (100 µl per well) overnight at 4 °C. After washing and blocking with 3% non-fat milk in PBS, purified Fabs were serially diluted threefold from 30 µg ml−1 and incubated for 1 h at room temperature. Bound Fab was detected using an anti-His primary antibody followed by an HRP-conjugated goat anti-mouse secondary antibody. TMB development was stopped after 15 min with 2 M sulfuric acid, and absorbance was read at 450 nm. Apparent EC50 values were obtained by fitting concentration-response curves in GraphPad Prism 8.

### Formation and purification of the IL-4Rα/Fab-16/anti-Fab nanobody complex

Purified Fab-16, IL-4Rα ECD, and anti-Fab nanobody (Thermo Fisher Scientific, catalogue 1033270500) were mixed at a 1:1:1 molar ratio and incubated for 2 h at 4 °C. The complex was purified by SEC (Increase S200, PBS; Supplementary Fig. 2), and fractions corresponding to the ternary assembly were pooled and concentrated for cryo-EM grid preparation.

### Cryo-EM grid preparation and data collection

UltraAuFoil R1.2/1.3 300-mesh holey grids were glow-discharged using a PELCO easiGlow system. Three microliters of complex were applied to each grid in a Vitrobot chamber maintained at 4 °C and 100% relative humidity. Grids were blotted for 3 s and plunge-frozen in liquid ethane.

Grids were screened on a 200-kV Glacios microscope and transferred to a 300-kV Titan Krios G4 equipped with a Falcon 4i direct electron detector and Selectris energy filter for automated data collection. Movies were recorded in super-resolution mode with a target defocus range of approximately −1.0 to −2.0 µm and a total dose of ∼50 e^-^/Å^2^. A total of 9,062 movies were collected.

### Cryo-EM data processing

Movies were processed in RELION 5 [6]. Beam-induced motion was corrected, and CTF parameters were estimated for each micrograph. After removal of micrographs with poor ice, contamination, drift, or unsatisfactory CTF fits, 7,540 micrographs were retained. Automated picking and extraction yielded 2,937,684 particles. Iterative 2D classification retained 1,708,018 particles, which were used for ab initio reconstruction and 3D classification. A well-resolved class containing 1,129,195 particles was selected for local 3D classification, 3D refinement, CTF refinement, and particle polishing. The final reconstruction was calculated from 173,323 particles and reached 2.74 Å resolution by the gold-standard FSC = 0.143 criterion (Supplementary Fig. 3 and Supplementary Table 2).

### Atomic model building and refinement

The IL-4Rα component of PDB 6WGL [23] was fitted into the cryo-EM density using UCSF Chimera [31]. The receptor and Fab models were adjusted manually in Coot [32] and refined in real space using REFMAC5 [33,34]. The model was evaluated using MolProbity [35].

## Structural analysis

Fab-IL-4Rα contacts were identified using PyMOL and UCSF Chimera. Backbone RMSD values were calculated after superposition of the IL-4Rα chain on the corresponding chain of PDB 6WGL using UCSF Chimera. Structural figures were prepared in UCSF Chimera and PyMOL.

## Data availability

The cryo-EM maps, half maps, mask, and atomic coordinates will be deposited in the Electron Microscopy Data Bank and Protein Data Bank. Fab sequences and construct details required to reproduce the study can be provided upon request, subject to completion of intellectual-property review.

## Code availability

No custom image-processing code was required for the current reconstruction.

## Acknowledgements

We thank our colleagues in the VIVA Biotech Cryo-EM Platform, Biologics R&D Platform, CADD & AIDD Platform, MARS Platform, Protein & Structure Platform, DEL Platform and Marketing Team for their experimental, computational, and project support, as well as for valuable discussions and insightful suggestions.

## Author contributions

X. L. conceived the study, supervised the development of the cryoFAST workflow and the cryo-EM studies, and drafted the manuscript. Y. L. performed the cryo-EM experiments and data analysis. S. D. and W. L. supervised antibody discovery, development, and production. J. D. performed antibody discovery and production, biochemical characterization, and cryo-EM sample preparation. Y. Q. supervised the AI-guided antibody design and optimization. L. L. performed the AI-guided antibody design and optimization. D. Q. supervised target protein production. All authors contributed to data interpretation, reviewed the manuscript, and approved the final version.

## Supplementary information

**Supplementary Figure 1.**
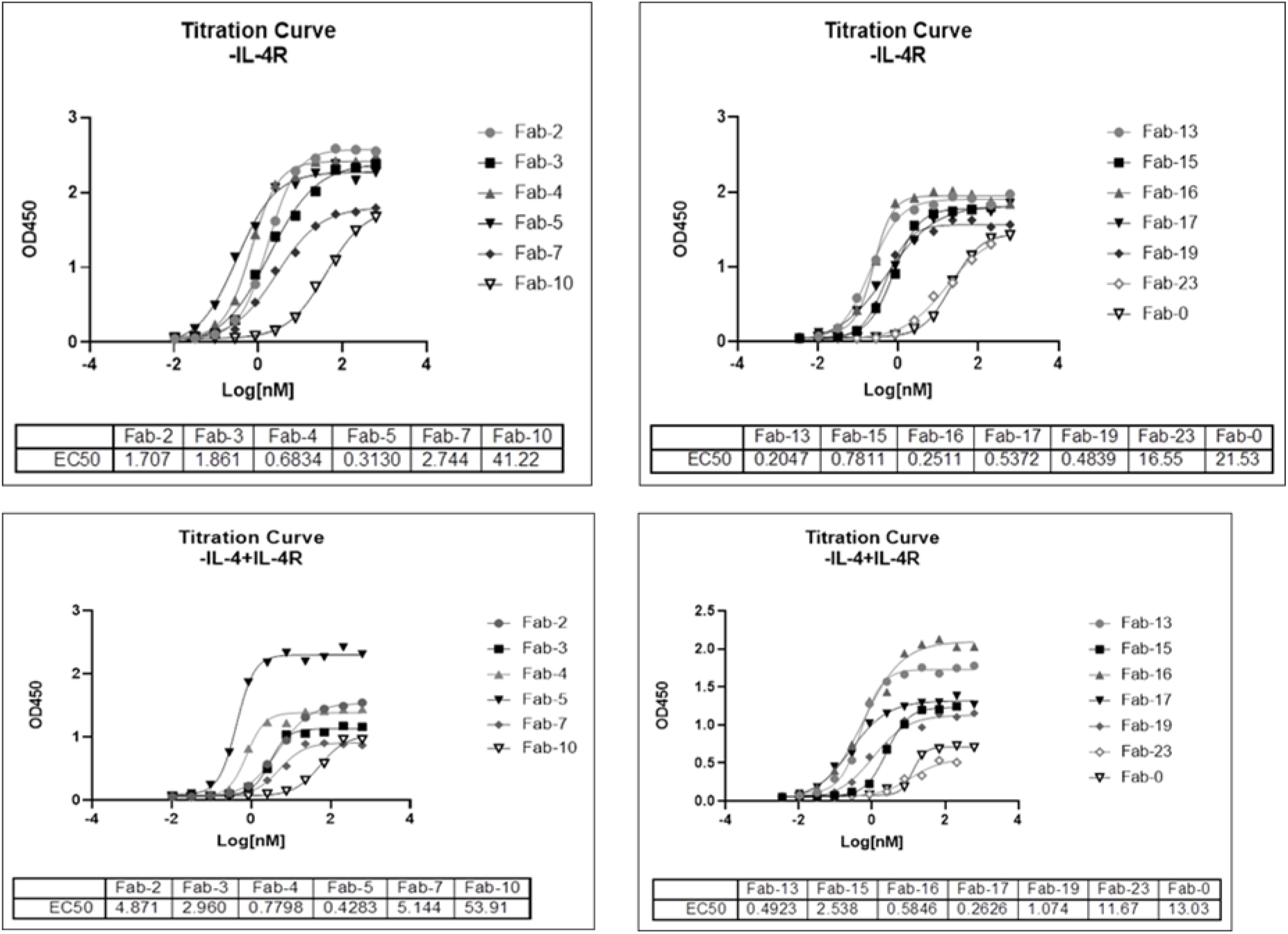
Full ELISA curves for Fab candidates. Concentration-response ELISA curves against the IL-4Rα ECD and a preassembled IL-4Rα-IL-4 complex are shown.

**Supplementary Table 1.** Apparent ELISA EC50 values for Fab candidates.

| Fab candidate | IL-4Rα ECD EC50 (nM) | IL-4Rα-IL-4 EC50 (nM) |
| --- | --- | --- |
| Fab-0 | 21.53 | 13.03 |
| Fab-2 | 1.707 | 4.871 |
| Fab-3 | 1.861 | 2.96 |
| Fab-4 | 0.6834 | 0.7798 |
| Fab-5 | 0.313 | 0.4283 |
| Fab-7 | 2.744 | 5.144 |
| Fab-10 | 41.22 | 53.91 |
| Fab-13 | 0.2047 | 0.4923 |
| Fab-15 | 0.7811 | 2.538 |
| Fab-16 | 0.2511 | 0.5846 |
| Fab-17 | 0.5372 | 0.2626 |
| Fab-19 | 0.4839 | 1.074 |
| Fab-23 | 16.55 | 11.67 |

**Supplementary Figure 2.**
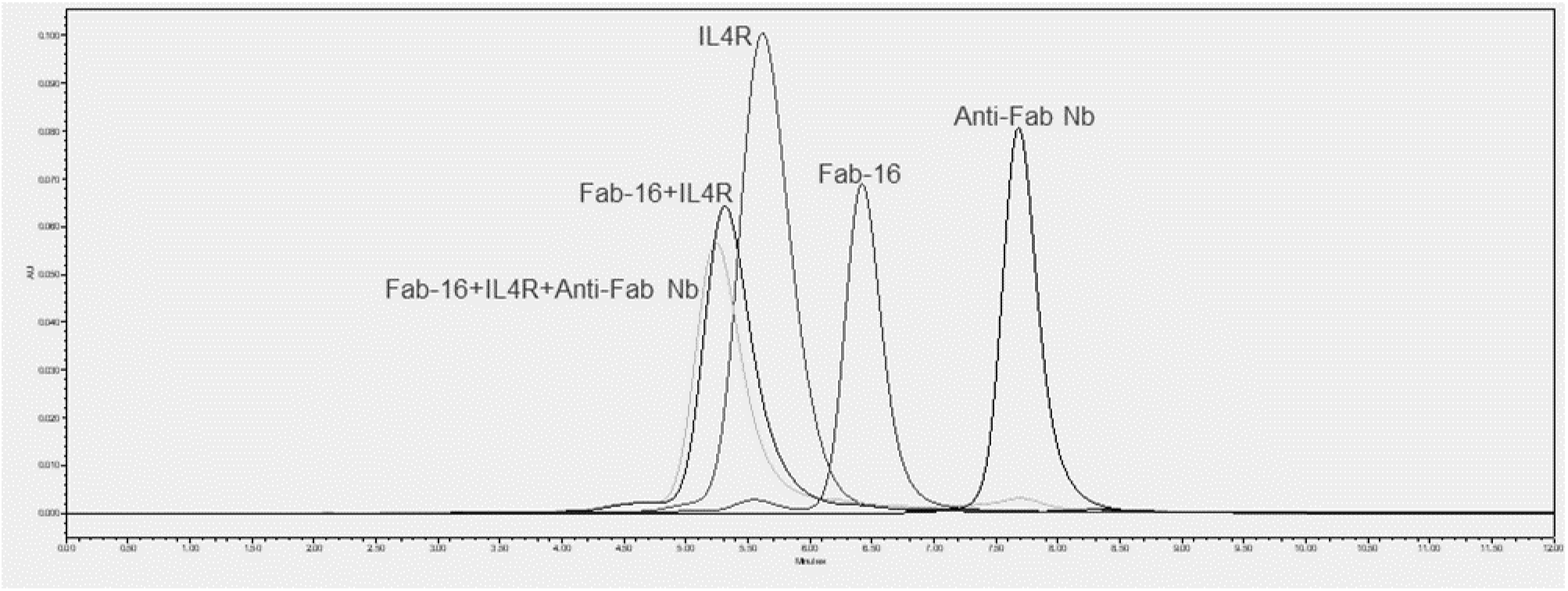
Analytical SEC of the IL-4Rα ECD, Fab-16, and anti-Fab nanobody alone and of the corresponding binary and ternary complexes.

**Supplementary Figure 3.**
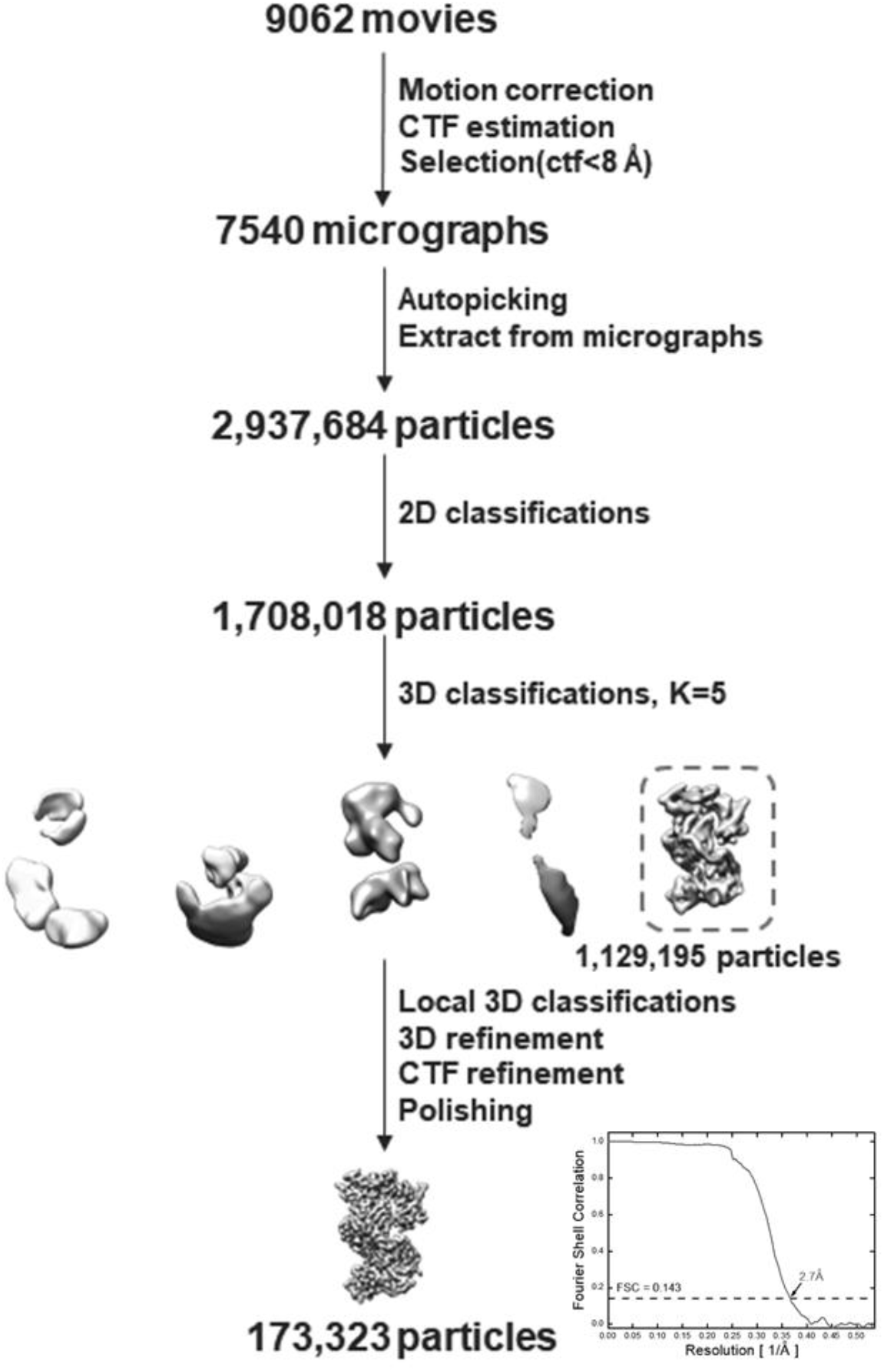
Cryo-EM processing workflow for the IL-4Rα ECD/Fab-16/anti-Fab nanobody complex.

**Supplementary Table 2.** Cryo-EM data-collection, processing, and refinement statistics.

| Parameter | Value |
| --- | --- |
| Microscope | Titan Krios G4 |
| Detector | Falcon 4i |
| Pixel size | 0.932 Å/pixel |
| Defocus | -1.0 to -2.0 μm |
| Movies | 9,062 |
| Final particles | 173,323 |
| Overall resolution | 2.74 Å (FSC = 0.143) |
| Ramachandran favored | 96.55 |
| Ramachandran outliers | 0.00 |
| Clashscore | 5.46 |
| MolProbity score | 1.52 |

